# The CYP27A1–VDR Feedback Rheostat Controls Mitochondrial Protein Homeostasis and Preserves Hippocampal-Striatal Network Integrity to rescue Cognitive impairment in HD condition

**DOI:** 10.64898/2026.06.11.731670

**Authors:** Vaishali Kumar, Pradeep Kodam, Chaitanya Kunja, Gayatri Brahmandam, Shreya N Balasubramanyam, Shashikant Patel, SKV Manjari, Sumana Chakravarty, Amartya Sanyal, Pragya Komal, Shuvadeep Maity

## Abstract

Emerging evidence links vitamin D (VD) deficiency to cognitive and motor dysfunction in Huntington’s disease (HD), yet the underlying mechanism remains unclear. Here, we combine *in vitro* genetic and 3-nitropropionic acid (3-NP)-induced *in vivo* models of HD to define the mechanistic basis of VD-mediated neuroprotection. We demonstrate that VD supplementation restores cognitive and motor deficits while improving mitochondrial function and cellular survival. Mechanistically, VD enhances mitochondrial fusion and rescues complex II expression, a key defect in HD pathology. We identify the mitochondrial enzyme CYP27A1 as a critical mediator of this effect, as its expression is reduced in HD models but restored upon VD treatment. Overexpression of CYP27A1 recapitulates the protective effects of VD, confirming its central role in maintaining mitochondrial integrity. Furthermore, VD promotes vitamin D receptor (VDR)-dependent transcriptional activation of CYP27A1 and additional nuclear- and mitochondrial-encoded genes, establishing a regulatory feedback loop that supports mitochondrial biogenesis and function. VD supplementation also improves proteostasis via attenuating endoplasmic reticulum stress. Together, our findings uncover a previously unrecognized VD–CYP27A1 axis that links mitochondrial dysfunction to HD pathology and highlight CYP27A1 as a potential therapeutic target.

## 1. Introduction

Huntington’s disease (HD) is a progressive neurodegenerative disorder characterized by motor and cognitive dysfunction, accompanied by the selective degeneration of striatal medium spiny neurons due to the accumulation of intracellular aggregates of mutant huntingtin (mHtt) protein^1^. These aggregates arise from the expansion of CAG trinucleotide repeats (>35) in the first exon of the huntingtin (HTT) gene, resulting in a toxic gain of function that disrupts multiple cellular processes, including mitochondrial dysfunction, ultimately leading to neuronal death. Postmortem brain studies, together with multiple *in vitro* and *in vivo* models of HD, have revealed alterations in the expression and activity of mitochondrial Complex II, resulting in bioenergetic failure and contributing to disease pathogenesis^2,3^.

Interestingly, clinical studies have indicated that HD patients with vitamin D (VD) deficiency or insufficiency exhibit exacerbated symptoms, including hyperkinetic movement and cognitive impairment^4–6^. Moreover, experimental studies involving induced alterations in VD status in various models further support the association between VD insufficiency and neurological disorders^7,8^. VD supplementation has been reported to improve motor dysfunction in HD and other neurodegenerative diseases ^9,10^. Furthermore, studies have shown that VD regulates mitochondrial function and redox homeostasis in the brain, suggesting a potential mechanistic link between VD signaling and HD pathophysiology ^11^.

While systemic vitamin D activation is well characterized, our research focuses on the local, mitochondrial-specific activation pathway. Within the brain, the presence of the Vitamin D Receptor (VDR) and vitamin D-metabolizing enzymes suggests the potential for local synthesis of active vitamin D within the central nervous system ^12^. Central to this process is CYP27A1, the only mitochondrial cytochrome P450 enzyme with dual catalytic activity involved in cholesterol metabolism and VD activation via its 25-hydroxylase activity^13,14^. Given that HD pathology is intrinsically linked to mitochondrial bioenergetic collapse, CYP27A1 is uniquely positioned to be a critical modulator of cellular health, yet its specific role in HD remains unstudied. CYP27A1 is the first 25-hydroxylase cloned and characterized in context to the VD metabolic pathway ^15^. Interestingly, CYP27A1 is the only mitochondrial dual enzyme balancing sterol and VD metabolism. While CYP27A1 also handles sterol clearance, our study focuses exclusively on its role as a mitochondrial VD 25-hydroxylase and its connection with the VDR mediated signaling.

In this study, we first demonstrated that vitamin D supplementation restores cognitive impairment and motor dysfunction in the 3-NP-induced HD mouse model, in which complex II inhibition induces HD-like behavioral symptoms. Interestingly, VD supplementation not only restores complex II expression but also enhances expression of mitochondrial CYP enzymes and VD receptors, which are involved in VD signaling, indicating a direct correlation between improved behavioral symptoms and mitochondrial health. Interestingly, *in vitro* genetic model of HD expressing toxic PolyQ(Q74) showed decreased expression of CYP27A1 and VDR, along with fragmented mitochondria in HD condition. Vitamin D (calcitriol) supplementation restored expression of both CYP27A1 and VDR with improved mitochondrial health and cellular survival. Further, VD supplementation induces transcriptional activation of nuclear and mt-DNA-encoded genes that promote mitochondrial biogenesis. Overexpression of CYP27A1 corroborates the effect of VD supplementation, confirming CYP27A1 as a local modulator of cellular and mitochondrial health in HD conditions. Our results reveal that CYP27A1-VDR feed-forward loop acts as a master regulator during VD supplementation. We provide evidence that at cellular level VD therapy not only restores mitochondrial bioenergetics by restoring Succinate Dehydrogenase Complex Iron Sulfur Subunit B (SDHB, Complex II) expression, normalization of CHCHD_4_ level and re-establishing functional mitochondrial dynamics but also promotes mitochondrial biogenesis by transcriptionally activating nuclear- and mt-DNA-encoded genes. In addition, it alleviates the proteostatic burden by attenuating ER stress markers and clearing mHtt aggregates. Importantly, restoration of behavioral and cognitive deficits in the HD mouse model following VD supplementation rescued CYP27A1 expression and mitochondrial health, confirming a novel role of CYP27A1 in functional recovery. By characterizing the CYP27A1-VDR axis, this study identifies a novel therapeutic framework for mitigating the bioenergetic failures that drive the progression of Huntington’s disease.

## 2. RESULTS

### 2.1. Vitamin D improves cognitive decline in the HD mice model

Both patients and animal model studies consistently reported spatial memory impairment in Huntington’s disease (HD)^16–18^. Individuals with HD exhibit marked deficits in spatial navigation, including difficulties in navigating environments, recalling spatial layouts, and recognizing objects or landmarks^19^. Although spatial memory has traditionally been associated with hippocampal dysfunction, recent studies demonstrated that disruption of effective connectivity between hippocampal and striatal regions was involved in spatial memory impairment in HD^20–23^. Thus, cognitive decline of HD patients potentially linked to functional collapse of hippocampal-striatal circuits.

To investigate potential neuroprotective strategies for spatial memory impairment in HD, we utilized a 3-nitropropionic acid (3-NP)-induced mouse model, a well-established pharmacological model that recapitulates key features of HD, including selective striatal degeneration, complex II dysfunction, and behavioral deficits. Spatial learning and memory were assessed using the established Morris water maze (MWM) protocol over four groups (n = 10/group) of mice: Control, HD, vitamin D (VD), and HD + VD (**Figure 1A**) ^23^. HD mice exhibited increased escape latency, reduced mean speed, and increased distance to target (path length), indicating impaired spatial learning, whereas VD supplementation (500 IU/kg, i.p., for 15 consecutive days) significantly improved these deficits (**Figure 1B, Supplementary Figure 1A**). In the probe trial conducted 24 hours later, VD-treated HD mice showed restored memory retention compared to untreated HD controls (**Figure 1B, Supplementary Figure 1A**). In addition, rotarod analysis showed that HD mice exhibited impaired motor coordination and reduced latency to fall, which was significantly improved following VD supplementation (**Supplementary Figure 1B**).

**Figure 1:**
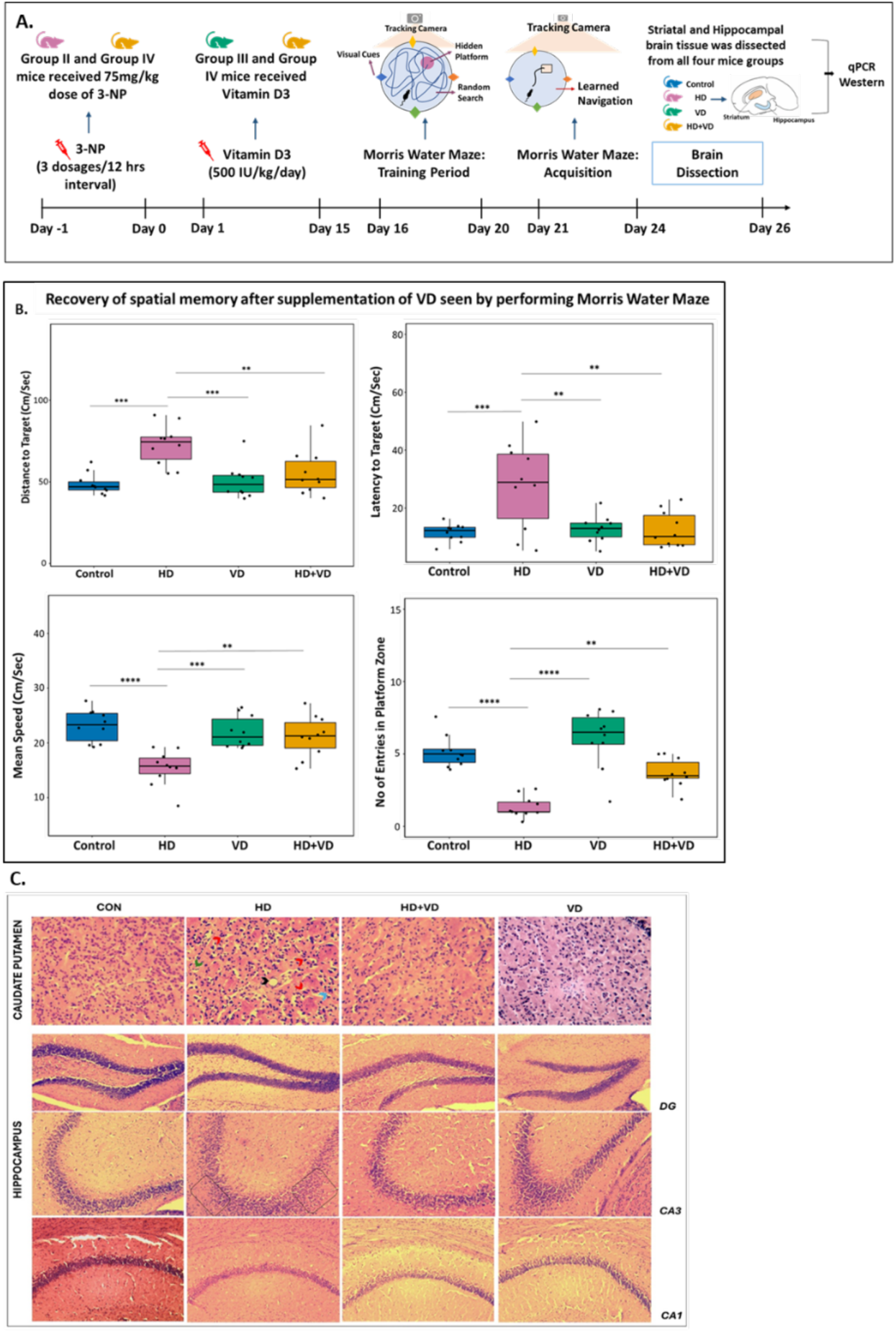
Vitamin D (VD) supplementation rescues cognitive deficits in 3-NP-induced mouse model of Huntington’s Disease (HD) (A) Schematic representation of the experimental design. Mice were administered 3-nitropropionic acid (3-NP) to induce HD-like pathology, followed by VD supplementation. Behavioral assessment was performed using the Morris water maze (MWM) to evaluate spatial learning and memory. After behavioral testing, striatum and hippocampus were collected for molecular analyses (qPCR and immunoblotting). (B) MWM performance showing the learning phase across successive training days (Panels 1-3) and probe trial assessing memory retention (Panel 4). VD supplementation significantly improved spatial learning and memory in 3-NP—treated mice (n = 10 per group). Quantified data represent three independent experiments (n = 3) and are presented as mean ± SD. Statistical significance was determined using unpaired Welch’s t-test with Holm’s correction (*p < 0.05, **p < 0.005, ***p < 0.0005, ****p < 0.00005). (C) Representative hematoxylin and eosin (H&E)-stained brain sections (20x magnification). Top row: caudate putamen (striatum); lower rows: hippocampal subregions (dentate gyrus [DG], CA3, and CA1). In HD mice, the striatum shows marked vacuolization (black arrows), pyknotic nuclei (red arrows), gliosis (green arrows), and degenerating cells (blue arrow). The CA3 region exhibits mild pyramidal layer disorganization (bracket). These pathological features are substantially ameliorated in VD-treated HD mice.

Following the behavioral assessment, we evaluated whether functional recovery correlated with structural changes in brain regions involved in spatial cognition. Histological analysis of the caudate-putamen (striatum) revealed pronounced neurodegeneration in HD mice compared with controls (**Figure 1C**). Control and VD-treated groups exhibited preserved striatal architecture with normal neuronal morphology (healthy, round neuronal nuclei and a uniform neuropil), whereas HD mice showed marked gliosis, vacuolization (spongiosis), and nuclear pyknosis, indicative of neurodegeneration. Notably, VD supplementation significantly attenuated these pathological features (**Figure 1C**).

In the hippocampus, the dentate gyrus (DG) and CA1 regions remained largely unaffected across groups; however, the CA3 region in HD mice showed mild disorganization of the pyramidal layer and focal neuronal loss (**Figure 1C**). These abnormalities were absent in VD-treated HD mice, which displayed preserved cytoarchitecture.

Collectively, these findings indicate that VD-mediated rescue of spatial memory deficits is associated with striatal neuroprotection and preservation of hippocampal integrity, thereby mitigating HD-associated neurodegeneration and sustaining cognitive function.

### 2.2. Vitamin D restores complex II (SDHB) dysfunction and mitigates neurodegeneration in Huntington’s Disease models

Post-mortem and experimental studies consistently demonstrate defects in oxidative phosphorylation in HD, including marked reduction in the expression as well as functionality in mitochondrial Complexes, specifically complex II^2,24,25^. Given that 3-NP, a potent and irreversible inhibitor of mitochondrial Complex II, recapitulates key aspects of HD-associated bioenergetic failure, we utilized the 3-NP model to investigate whether VD could modulate Complex II integrity^26,27^. *In vivo*, 3-NP administration resulted in a significant downregulation of Complex II components in the striatum and hippocampus at both transcript and protein levels. Notably, regulated VD supplementation restored the expression of complex II components (SDHA & SDHB), thereby indicating recovery of Complex II expression (**Figure 2A, B & Supplementary Figure 2A**). These molecular alterations likely underlie the observed improvement in behavioral deficits, highlighting the functional significance of Complex II restoration. Consistently, in an *in vitro* HD model expressing mutant Q74 huntingtin, similarly Complex II components (SDHA & SDHB) were found to be upregulated upon VD treatment, recapitulating the *in vivo* findings (**Figure 2C–E, Supplementary Figure 2B, C**). Furthermore, functional assessment by flow cytometry revealed a marked reduction in viable cells under HD conditions, which was significantly rescued upon VD treatment (**Figure 2F**), supporting its cytoprotective role. Collectively, the restoration of Complex II expression and associated recovery of cell viability establishes a mechanistic framework for VD–mediated neuroprotection in HD models.

**Figure 2:**
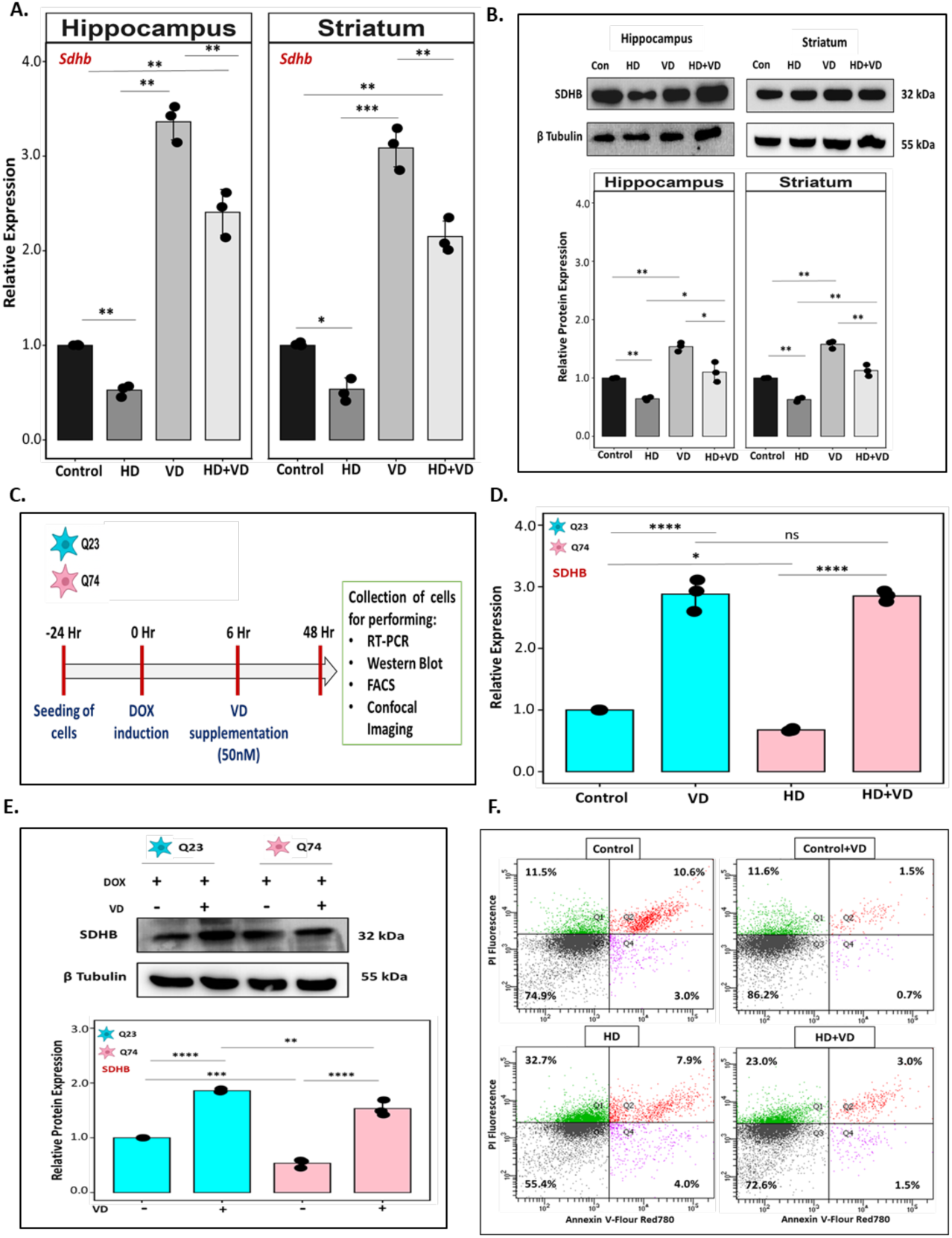
Vitamin D supplementation restores the expression of mitochondrial Complex II subunit in HD models and improves cellular viability. (A, D) RT-qPCR analysis shows a significant reduction in SDHB (a key subunit of mitochondrial Complex ll/succinate dehydrogenase) under HD conditions, indicating impaired Complex II integrity. VD supplementation significantly restores SDHB transcript levels (n = 3). (C) Schematic represents experimental outline on VD supplementation to *in vitro* lines expressing PolyQ. repeats (Q23/Q74). (B, E) Representative western blots confirm restoration of SDHB expression in HD condition upon VD treatment (n = 3). (F) Flowcytometric analysis (n=3) showed VD supplementation improves cellular viability. The dot plots of different experimental conditions (Control, VD, HD, and HD+VD) represents four quadrants Q1 (Top Left): Necrotic or debris population (Annexin V negative / PI positive), Q2 (Top Right): Late apoptotic or necrotic cells (Annexin V positive / PI positive), Q3 (Bottom Left): Live cells (Annexin V negative / PI negative) and Q4 (Bottom Right): Early apoptotic cells (Annexin V positive / PI negative). Data are presented as mean ± SD from three independent experiments. Statistical significance was determined using unpaired Welch’s t-test with Holm’s correction: P < 0.05, *P < 0.005, **P < 0.0005, and ***P < 0.00005.

### 2.3. Vitamin D restores mitochondrial function in HD Models

Earlier postmortem analysis of the HD striatum by Kim et al. (2010) demonstrated significant alterations in mitochondrial dynamics. These were characterized by increased Drp1-mediated fission and reduced Mfn1-mediated fusion, alongside pronounced bioenergetic deficits that contribute to neuronal degeneration^28^. Building on evidence of complex II recovery, we systematically evaluated the effects of VD on parameters linked to mitochondrial health, including mitochondrial dynamics, reactive oxygen species (ROS) production, membrane potential (Ψm), and the expression patterns of fission–fusion regulators.

Using both *in vivo* and *in vitro* genetic models of HD, we confirmed a significant downregulation of fusion genes (*MFN1/2*) and an upregulation of fission genes (*DRP1* and *MFF*) under HD conditions (Figure 3A, Supplementary Figure 3B-D). These findings align with previous reports showing that mutant huntingtin (mHtt) interacts with DRP1, enhancing its activity and promoting mitochondrial fragmentation^29^. Notably, VD treatment restored the fission–fusion balance in HD (Q74) cells by increasing *MFN1* expression while reducing *DRP1* and *MFF* levels (Supplementary Figure 3D). Corroborating these *in vitro* findings, the 3-NP mouse model exhibited enhanced fission and reduced fusion gene expression in both the hippocampus and striatum. VD supplementation reversed these alterations, restoring fusion gene expression and reducing fission markers (**Figure 3A; Supplementary Figure 3B-D**).

**Figure 3:**
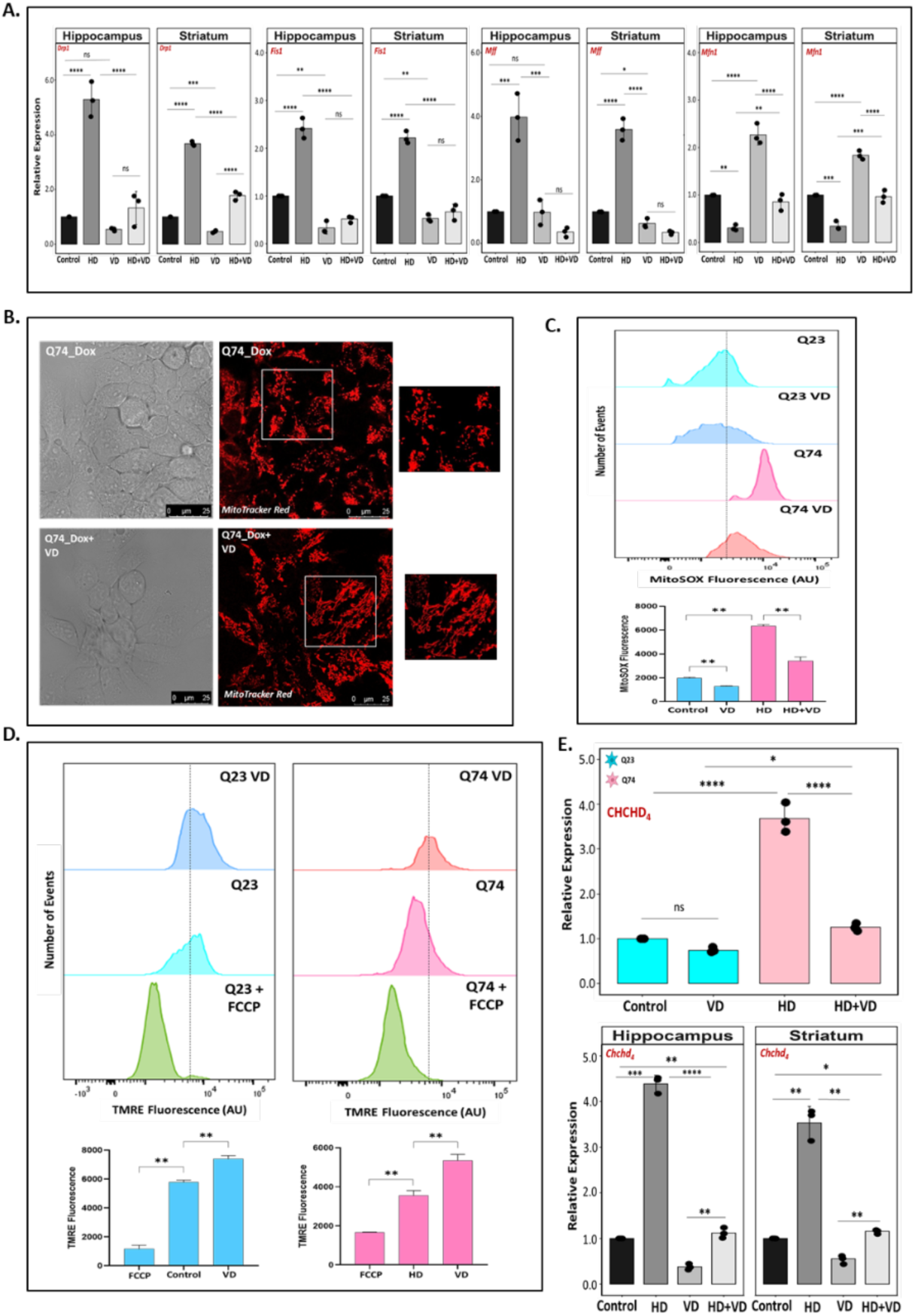
Vitamin D supplementation restores mitochondrial morphology, membrane potential, and suppresses ROS production in HD condition. (A) HD Condition shows higher expression of fission genes (DRP1, FIS1 and MFF) and reduced expression of fusion gene (MFN1) in the brain regions indicating mitochondrial fragmentation (n=3). Supplementation of VD significantly increases fusion gene expression whereas reduces fission gene expression indicating restoration of mitochondrial dynamics. (B) Confocal images of mitochondria stained with MitroTracker Red shows severe mitochondrial fragmentation in *in vitro* model of HD and VD supplementation restores elongated mitochondrial network. (C) Representative flow cytometry analysis reveals elevated mitochondrial ROS in HD condition. VD supplementation decreases mitochondrial ROS (n=3). (D) TMRE-based quantification indicates loss of mitochondrial membrane potential (Δψm) in HD (Q74) cells while VD treatment restores the membrane potential in the HD cells (n=3). (E) RT-PCR data shows significantly elevated expression of CHCHD_4_ in HD condition which upon VD treatment reduced to basal level as observed in control condition. Bar plot is from three independent experiments (n = 3) and all values are presented as mean with SD; *p value calculated using unpaired Welch’s t-test with Holm’s correction *P < 0.05. *p < 0.05, **p < 0.005, ***p < 0.0005, and ****p < 0.00005).

Furthermore, confocal imaging using MitoTracker Red revealed that HD cells displayed fragmented, punctate mitochondria with reduced mitochondrial footprint, whereas VD treatment restored elongated, tubular mitochondrial networks and significantly increased mitochondrial footprint, indicative of enhanced mitochondrial fusion (**Figure 3B; Supplementary Figure 3A**). Fragmented mitochondria exhibit reduced electron transport efficiency, leading to increased electron leakage and directly causing the overproduction of ROS. Analyses of post-mortem HD brains have demonstrated significantly elevated oxidative damage, including increased markers of lipid peroxidation, protein oxidation, and DNA oxidation in the striatum and cortex. Concordant findings from transgenic mouse models and neuronal HD cell systems consistently show enhanced ROS production and redox imbalance, establishing oxidative stress as a major pathogenic driver in HD^30–33^. Comparable pathogenic conditions were recapitulated in our *in vitro* genetic model of HD, in which mitochondrial ROS were significantly elevated, as assessed with the mitochondrial superoxide–sensitive fluorophore MitoSOX Red (**Figure 3C)**. VD– induced enhancement of mitochondrial fusion restored mitochondrial homeostasis, which was accompanied by a marked reduction in mitochondrial ROS and an overall improvement in mitochondrial health (**Figure 3C**).

In addition, several studies in HD patients and experimental models also reported reduced mitochondrial membrane potential (ΔΨm) and increased susceptibility to Ca²⁺-induced mitochondrial permeability transition, indicating bioenergetic stress associated with reduced ATP production and compromised mitochondrial function^29,34,35^. Notably, mitochondrial membrane potential (Δψm) loss is closely linked to increased oxidative stress. In our *in vitro* model, VD supplementation restored mitochondrial membrane potential, indicating a healthier bioenergetic state (**Figure 3D**). Overall, our study demonstrates that VD acts as a potent neuroprotective agent that mitigates the core bioenergetic failures and oxidative stress hallmark of Huntington’s Disease (HD) pathology. Mechanistically, VD promotes mitochondrial fusion, enhances mitochondrial metabolism by restoring membrane potential, and reduces reactive oxygen species (ROS), ultimately preventing mitochondrial fragmentation under HD pathological conditions.

Interestingly, our investigation revealed significant alterations in CHCHD_4_ (the mammalian homolog of yeast Mia40), a critical protein involved in mitochondrial protein import and redox regulation. Both gene and protein expression analyses showed that CHCHD_4_ levels were significantly elevated in our *in vivo* and *in vitro* HD models. This aligns with prior literatures while Schlagowski et al. demonstrated that increased levels of the mitochondrial import factor Mia40 could prevent polyQ protein aggregation and augment cytosolic proteostasis, a subsequent study on HD proposed that the upregulation of CHCHD family proteins serves as a compensatory protective mechanism against disease-induced oxidative stress^36,37^.Notably, we observed a reduction in CHCHD_4_ expression following VD supplementation. Rather than merely suppressing the CHCHD_4_ response, VD treatment effectively normalizes this previously strained compensatory mechanism. This restoration of homeostatic balance is further evidenced by rescued mitochondrial dynamics, stabilized membrane potential, and reduced ROS production (**Figure 3E, Supplementary Figure 3C**). By modulating the fission-fusion equilibrium and reinforcing the mitochondrial network, VD supplementation ultimately restores cellular proteostasis and alleviates HD-induced mitochondrial dysfunction.

### 2.4. Vitamin D metabolic pathway: Upregulation of CYP27A1 and VDR as a mechanism for neuroprotection

VD is increasingly recognized as a neuroactive hormone, and its active form binds to the vitamin D receptor (VDR) to initiate downstream signaling events. VDR is a ligand-activated transcription factor widely expressed both in neurons and glial cells across key brain regions, including the hippocampus, cortex, and basal ganglia, where it regulates gene expression associated with neuronal survival, neurotrophic support, and cellular homeostasis^38,39^. Crucially, the brain possesses autonomous enzymatic machinery, consisting of the hydroxylases CYP27A1, CYP2R1, CYP27B1, and the catabolic enzyme CYP24A1, to facilitate the local synthesis and availability of active VD, thereby supporting a self-sustaining intracrine system within the central nervous system^40–43^ (Figure 4A). Despite the functional necessity of this local system, clinical data consistently reveal a high prevalence of VD deficiency among patients with Huntington’s Disease (HD)^6,44,45^

**Figure 4:**
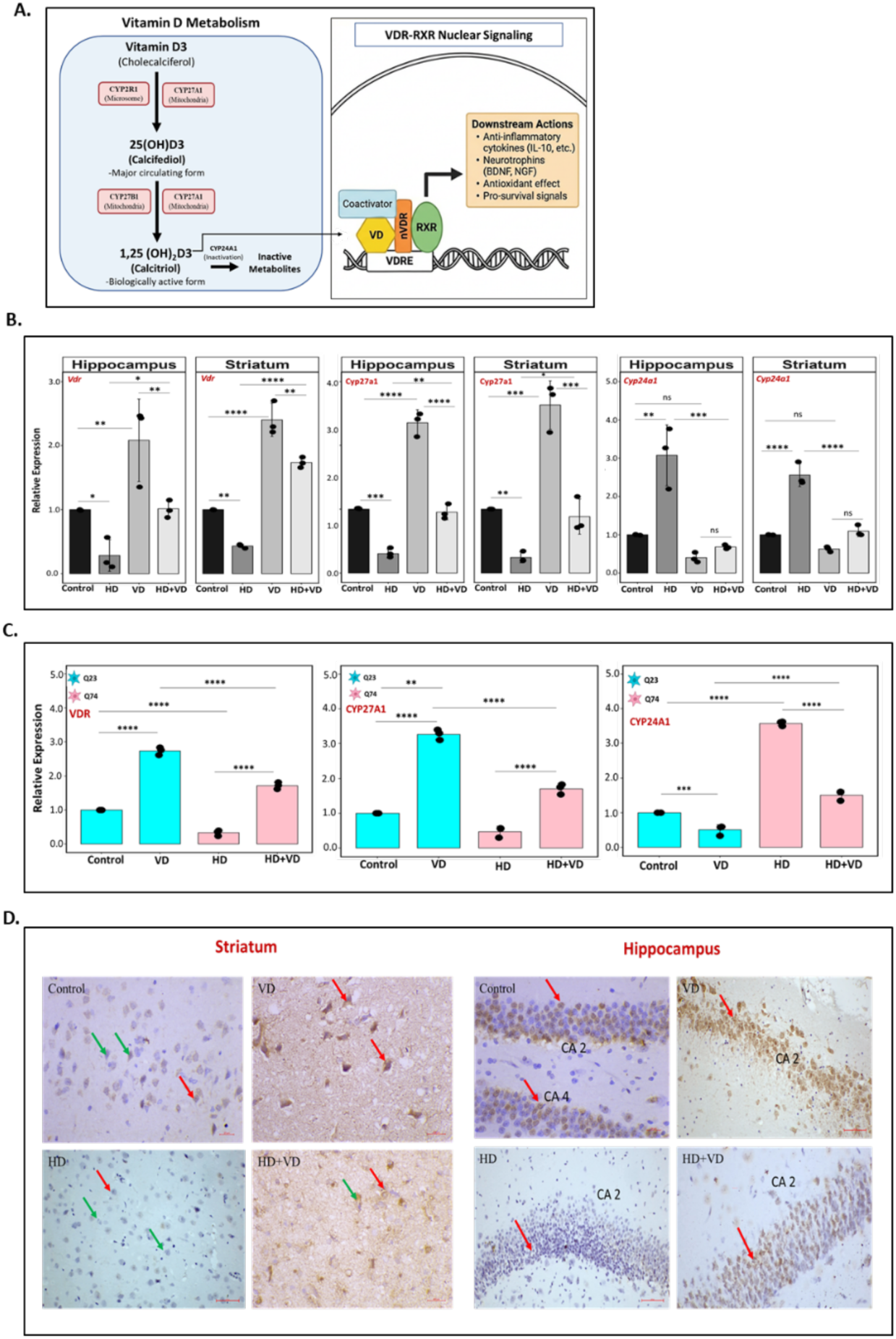
Dysregulation of the Vitamin D metabolic axis in in vivo and in vitro HD models. (A) Schematic representation of the VD pathway. (B) The HD group shows a significant downregulation of VDR and CYP27A1 expression in both the brain regions compared to control, whereas VD supplementation significantly restores its expression. In contrast, CYP24A1 expression was significantly elevated in HD condition and normalized upon VD treatment (n=3). (C) Gene expression analysis of VDR, CYP27A1, and CYP24A1 in the in vitro HD model confirms in vivo findings (n=3). (D) Representative images of VDR immunohistochemistry of the striatum and hippocampus shows reduced VDR immunoreactivity in the HD condition (IHC score = 0, 1), while VD supplementation recovers VDR expression (HD+VD; IHC score = 2, 2). (VDR-positive neurons and glial cells identified by blackish-brown staining, whereas VDR-negative cells appear blue due to counterstaining. Red arrows indicate VDR-positive cells, whereas green arrows indicate weakly stained or negative cells). Bar plot is from three independent experiments (n = 3) and all values are presented as mean with SD; *p value calculated with using unpaired Welch’s t-test Holm’s correction *P < 0.05. *p < 0.05, **p < 0.005, ***p < 0.0005, and 44D < 0,00005).

Aligning with these clinical observations, our previous work demonstrated that VDR expression is significantly downregulated in HD experimental models^12,46,47^. In the present study, we investigated the underlying molecular mechanisms driving this deficiency by mapping the expression patterns of the major mitochondrial cytochrome P450 (CYP) enzymes involved in the VD metabolic pathway. Notably, we identify mitochondrial CYP27A1 downregulation—a dual-action resident enzyme capable of regulating two consecutive hydroxylation steps required to generate active calcitriol—as a novel molecular signature of HD pathology. It is likely a primary driver of both localized and systemic reductions of active VD production. Quantitative RT-qPCR analyses demonstrated a significant decrease in the expression of CYP27A1 under HD conditions (Figure 4B–C). In contrast, analysis of the alternative hydroxylating enzyme, CYP2R1, showed no significant alterations (Supplementary Figure 4C), while the active VD-degrading enzyme, CYP24A1, was significantly upregulated in the HD state, establishing a hyper-catabolic environment that further depletes active ligand (VD) availability (Figure 4B–C). The VD supplementation restored the expression of CYP27A1 while normalizing the aberrantly high basal levels of the catabolic enzyme CYP24A1 enhancing the tissue’s intrinsic capacity to synthesize and maintained active VD levels (Figure 4B–C).

Importantly, therapeutic supplementation with VD successfully reversed these pathological deficits in both our *in vivo* and *in vitro* HD models. Quantitative PCR and western blotting confirmed that VD supplementation significantly restored VDR expression (Figure 4B-C, Supplementary figure A-B). Immunohistochemical analyses showed moderate basal VDR immunoreactivity in the control brain regions. In the HD group, VDR staining was markedly reduced (fewer brown-positive cells and overall reduced signal) in both the striatum and hippocampus indicating loss of VDR expression under diseased conditions. In contrast, VD supplementation increased VDR immunoreactivity (increased overall brown staining) in the HD+VD in both regions relative to untreated HD tissue (**Figure 4D**).

Taken together, these findings demonstrate that VD treatment does not merely replace a missing nutrient; rather, it effectively re-establishes localised intrinsic metabolic capacity, upregulates VDR availability, and ensures a neuroprotective feedback mechanism capable of mitigating HD-induced neurodegeneration.

### 2.5. Vitamin D promotes transcriptional activation of nuclear and mt-DNA encoded genes regulating mitochondrial function

Given that both our *in vivo* and *in vitro* models exhibited severe mitochondrial dysfunction—a well-established hallmark of HD)—we sought to determine whether VD supplementation could restore overall mitochondrial health and integrity. Prior studies in non-neuronal systems demonstrated that VDR-mediated signaling can activate downstream pathways governing mitochondrial biogenesis ^48–51^. Driven by this premise, we investigated whether the therapeutic restoration of VDR expression could activate mitochondrial biogenesis pathways under HD conditions. Mitochondrial biogenesis requires coordinated nuclear and mitochondrial gene expression driven by the PGC-1α–NRF1–TFAM axis, where NRF1 binds the TFAM promoter to regulate mitochondrial DNA (mtDNA) replication. Although impaired PGC-1α function routinely underlies bioenergetic failure in HD, PGC-1α can directly serve as a transcriptional coactivator for the VDR itself^52–57^, structurally linking VD signaling to mitochondrial gene regulation. Consistent with this framework, our *in vitro* HD model exhibited increased expression of PGC-1α, NRF1, and TFAM following VD supplementation, paralleling the restoration of VDR levels (**Figure 5A, Supplementary Figure 5A**). This transcriptional activation was accompanied by a significant increase in mtDNA copy number, in agreement with elevated TFAM expression (**Figure 5B**). Furthermore, VD treatment enhanced the expression of COX1 (a mitochondria-encoded gene) and COX4 (a nuclear-encoded component of the electron transport chain), mirroring the observed upregulation of the complex II gene SDHB (**Figure 5C, Supplementary Figure 5A, B**). Collectively, these findings demonstrate that VD activates a coordinated mito-nuclear transcriptional program in HD, characterized by increased mtDNA content and enhanced expression of respiratory chain genes. By engaging the VDR–PGC-1α–NRF1–TFAM signaling cascade (**Figure 5D**), VD not only improves mitochondrial physiology but also restores coordinated genomic communication, which is essential for cellular bioenergetics in the context of HD pathology.

**Figure 5:**
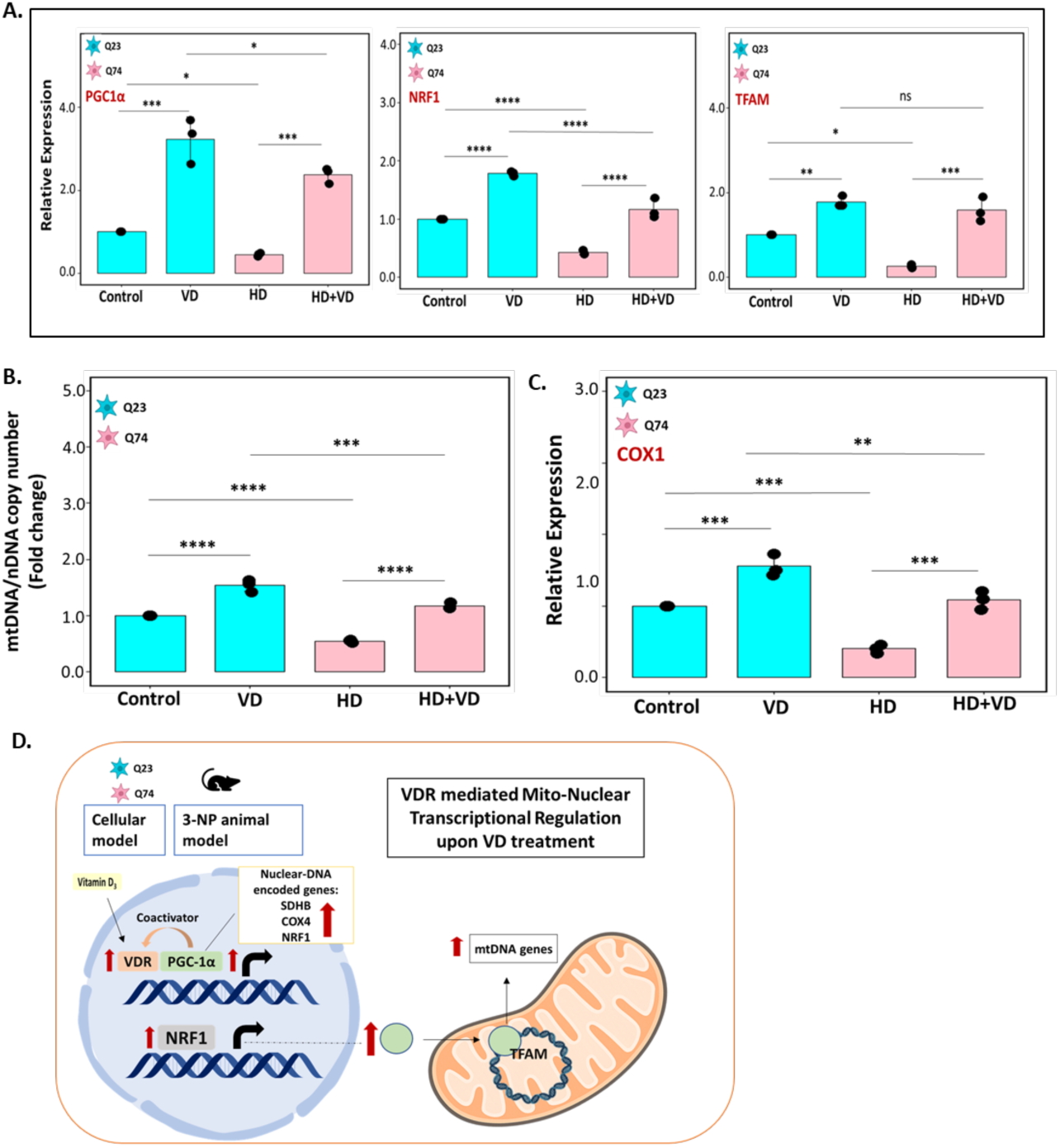
Vitamin D activates the VDR-PGC-1a-NRF1-TFAM axis to restore mitochondrial gene expression. (A) Relative mRNA expression shows significant downregulation of PGC-1a, NRF1, and TFAM in HD conditions, which is restored upon VD supplementation (n=3). (B) qPCR confirms increased mitochondrial DNA (mtDNA) copy number after VD treatment (n=3). (C) Mitochondrial-encoded respiratory gene COX1 is reduced in HD condition, which is significantly increased upon VD treatment (n=3). (D) Schematic representation illustrating nuclear and mitochondrial encoded gene expression after VD treatment. Bar plot is from three independent experiments (n = 3) and all values are presented as mean with SD; *p value calculated using unpaired Welch’s t-test with Holm’s correction *P < 0.05. *p <0.05, **p <0.005, ***p < 0.0005, and ****p < 0.00005).

### 2.6. CYP27A1-VDR feed-forward loop regulates mitochondrial proteostasis and cellular survival in HD

To determine whether reduced CYP27A1 directly contributes to mitochondrial dysfunction in HD—rather than merely impairing local calcitriol synthesis—we investigated whether overexpressing this inner membrane-resident enzyme could bypass the observed pathological deficits and rescue network dynamics. In both control and HD lines, CYP27A1 overexpression significantly upregulated VDR expression, effectively reconstituting the proximal signaling node of the localized Vitamin D pathway (**Figure 6A, 6B**).

**Figure 6:**
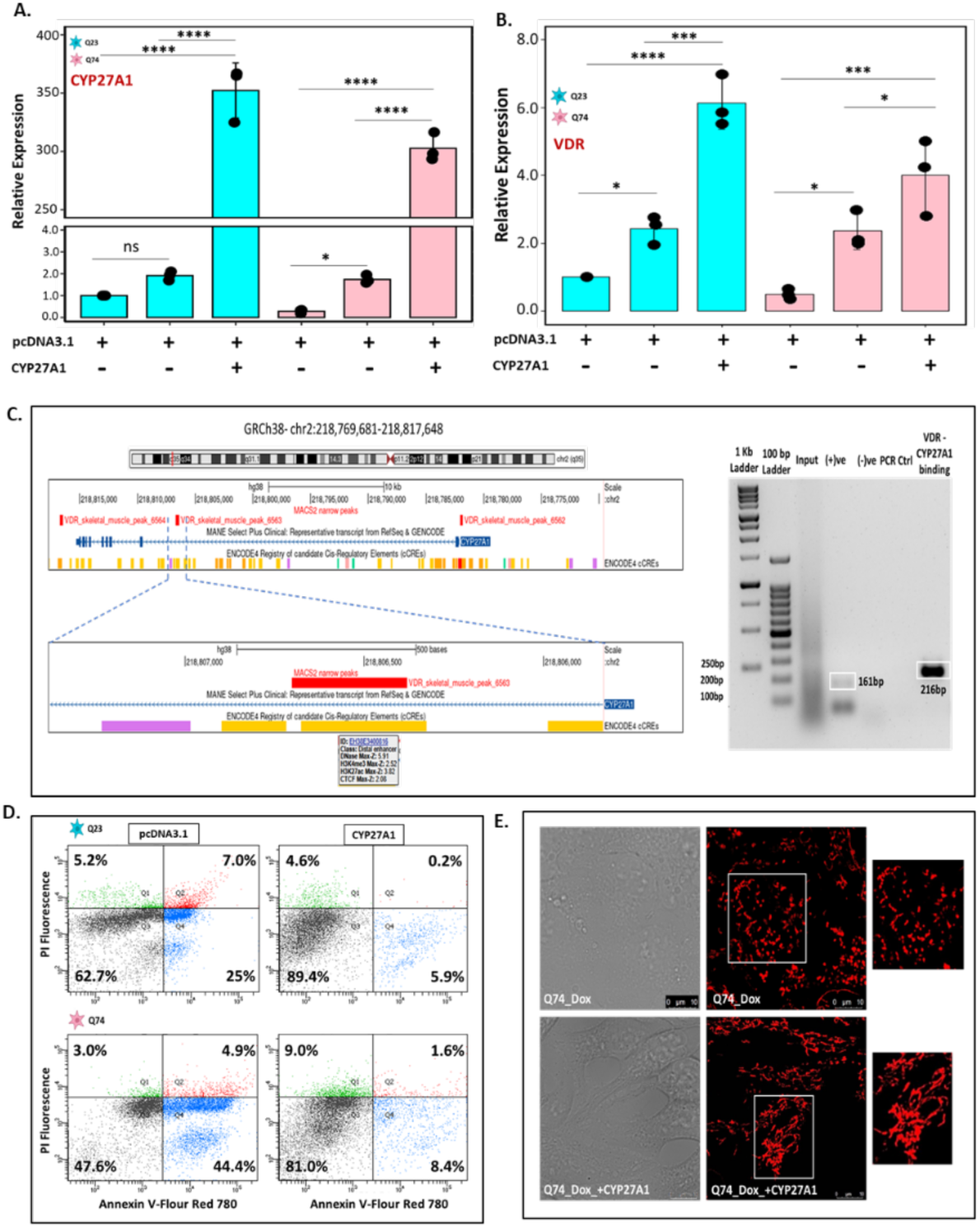
CYP27A1 overexpression enhances VDR expression with subsequent restoration of mitochondrial dynamics with increased cellular viability in *in vitro* HD model. (A, B) CYP27A1 overexpression significantly elevates CYP27A1 and VDR expression indicating activation of endogenous VD signaling (n=3). (C) UCSC genome browser snapshot (hg38 chr2:218,769,681-218,817,648) showing three VDR peaks (MACS2) at the CYP27A1 gene locus. The zoomed-in intronic VDR peak (6563) overlaps a distal enhancer region predicted by the ENCODE4 Registry of candidate cis-regulatory elements. ChIP-PCR with 2 independent biological replicates confirmed the binding of VDR at the predicted binding site (peak 6563) of CYP27A1 region. (D) Flowcytometric analysis of HD cells exhibits increased apoptotic population compared to control cells, whereas CYP27A1 overexpression significantly reduces both early and late apoptotic cell population, indicating improved cellular survival. (E) Confocal imaging shows restoration of elongated mitochondria in HD cells after CYP27A1 overexpression. Bar plot is from three independent experiments (n = 3) and all values are presented as mean with SD; *p value calculated using unpaired Welch’s t-test with Holm’s correction *P < 0.05. *p < 0.05, **p < 0.005, ***p < 0.0005, and ****p < 0.00005).

Remarkably, this genetic metabolic intervention completely mimicked the therapeutic effects of exogenous VD supplementation by triggering a robust downstream mitochondrial biogenesis program. Specifically, CYP27A1 induction drove a marked increase in the expression of the master transcriptional coactivator PGC-1α and its downstream effector TFAM (Supplementary **Figure 6A**), demonstrating that restoring this single mitochondrial enzyme is sufficient to reactivate the compromised mito-nuclear transcriptional axis in HD pathology. This transcriptional activation was accompanied by a marked elevation in both nuclear-encoded (SDHB, SDHA) and mitochondria-encoded (COX1) respiratory chain subunits, demonstrating a coordinated restoration of the mito-nuclear transcriptional program (**Supplementary Figure 6B, C)**. Crucially, this genetic rescue translated into structurally improved mitochondrial fusion dynamics (**Figure 6E**). This effect closely mirrored our findings with pharmacological intervention. CYP27A1 overexpression successfully reversed the aberrant expression profile of fission-fusion genes typically observed in the HD state, thereby restoring balanced mitochondrial dynamics.

Restoration of mitochondrial health prompted us to monitor cellular resilience after CYP27A1 overexpression. Flow cytometric analysis using Annexin V and Propidium Iodide (PI) staining revealed that HD-associated apoptosis was significantly attenuated following CYP27A1 overexpression. In HD cells, the proportion of surviving (Annexin V −ve/PI −ve) cells increased from approximately 47.6% to 81.0% upon CYP27A1 induction, with a concomitant reduction in late apoptotic and necrotic populations (**Figure 6D**).

Interestingly, we observed that CYP27A1 overexpression increased VDR expression, which is positioned downstream of the CYP27A1 metabolic pathway. Conversely, we observed increased VDR expression when the HD *in vitro* model was supplemented with exogenous active VD, calcitriol. These findings indicate the presence of a feedback loop balancing the CYP27A1–VDR–PGC-1α–NRF1–TFAM expression axis. To determine if this regulation occurs directly, we investigated whether a VDR binding site is present in the CYP27A1 gene. By analyzing publicly available chromatin immunoprecipitation sequencing (ChIP-Seq) data (GSE243777) of VDR binding in human skeletal muscle after anterior cruciate ligament reconstruction surgery ^58^, we identified three VDR binding sites at the CYP27A1 locus. One site (peak 6562) coincided with the CYP27A1 promoter, while the other two sites were located within an intronic region (peak 6563) and a 3′ downstream region (peak 6564). Both distal VDR binding sites overlapped with putative enhancer elements annotated in the ENCODE4 Registry of Candidate Cis-Regulatory Elements (cCREs) (Figure 6C, Supplementary file - ChIP peaks). ChIP-PCR analysis of one of the representative sites (peak 6563) confirmed VDR binding to CYP27A1 (**Figure 6C**). Given that CYP27A1 functions as a VD 25-hydroxylase, its upregulation likely enhances the local conversion of VD intermediates, thereby sustaining VDR activation. This positive feedback loop suggests that CYP27A1 not only initiates VD signaling but also acts as a rheostat to maintain intracellular metabolic competence under pathological stress. Collectively, our data reveal a novel regulatory feedback mechanism linking VD metabolism to receptor availability.

These results identify mitochondrial CYP27A1 as a critical neuroprotective factor in HD. By orchestrating a VDR-dependent feedback loop and activating the core PGC-1α–NRF1–TFAM biogenesis pathway, the targeted restoration of CYP27A1 effectively reverses bioenergetic failure, rescues structural network dynamics, and prevents the accelerated cell death characteristic of HD pathology. Crucially, our study identifies mitochondrial CYP27A1 downregulation, for the first time, as a key molecular signature of HD pathology. By linking the localized loss of this inner membrane-resident enzyme directly to the breakdown of mito-nuclear communication, these findings establish that repairing intrinsic metabolic machinery—rather than merely replacing downstream ligands—is a viable and potent therapeutic strategy to counteract neurodegeneration.

### 2.7. Vitamin D attenuates ER Stress–Driven proteostasis burden in Huntington’s Disease Models

VD has recently emerged as an important regulator of proteostasis, with accumulating evidence demonstrating its ability to suppress ER stress signaling across diverse biological systems and neurodegenerative disease models^59,60^. Neurodegenerative disorders are broadly characterized by the accumulation of misfolded proteins that form amyloid-like aggregates enriched in β-sheet structures, a hallmark observed across diseases such as Alzheimer’s disease (AD), Parkinson’s disease (PD), amyotrophic lateral sclerosis (ALS), and HD^61,62^. The persistent accumulation of misfolded proteins inside the ER can trigger ER stress^63,64^. In addition, impairment of ER-associated degradation (ERAD) can activate ER stress, as observed after accumulation of polyglutamine aggregates in the cytoplasm^65^. Based on these considerations, we also examined the expression of key ER stress markers following VD supplementation to evaluate its impact on proteostasis under disease conditions^66^.

Consistent with this, our findings demonstrate that VD supplementation reduced aggregate burden and suppressed ER stress in *in vitro* HD models. In our cellular system, aggregate load was assessed by GFP western blot, and in the HD condition, we observed enhanced GFP expression in insoluble protein fractions (aggregates) compared to the control. Notably, VD treatment led to a marked reduction in GFP-positive aggregates in the HD condition (**Supplementary Figure 7**). Furthermore, assessment of ER stress markers by RT–qPCR revealed significant attenuation of GRP78, PERK, ATF4, and sXBP1 (**Figure 7**). These data indicated VD mediated alleviation of ER stress and mutant huntingtin–induced proteotoxicity.

**Figure 7:**
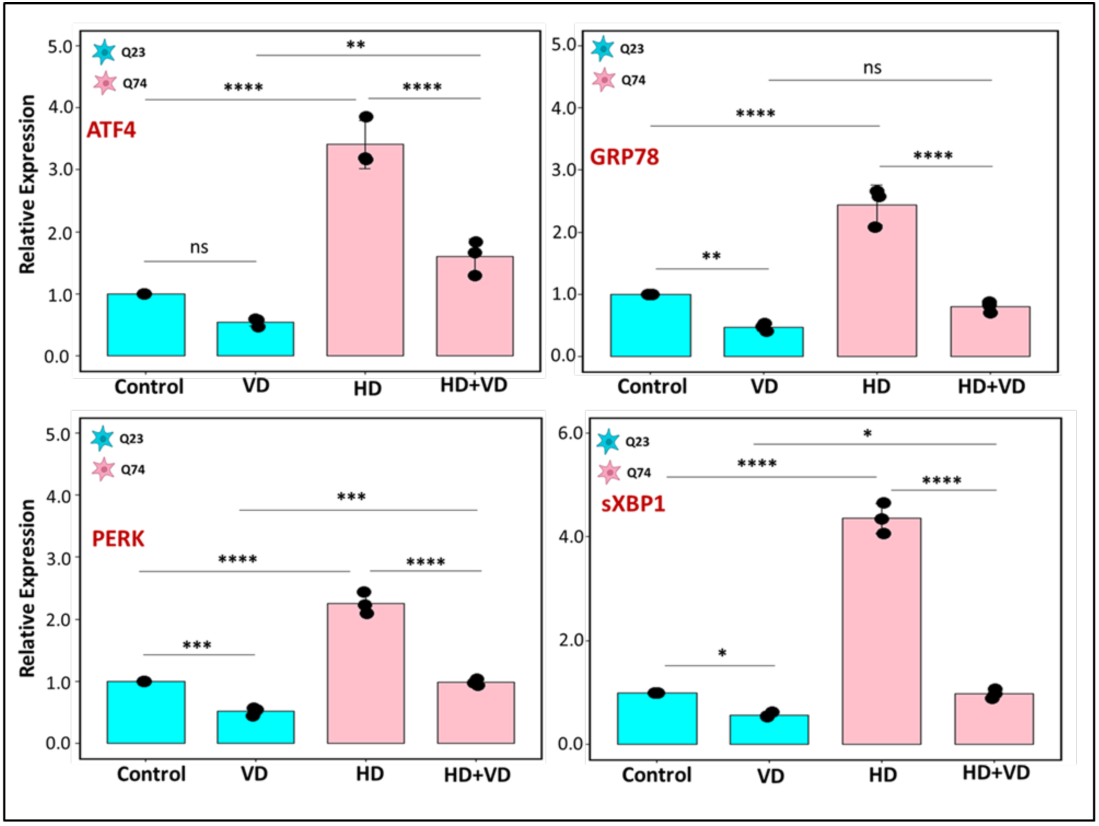
Vitamin D attenuates ER stress marker expression in Huntington’s disease cell model. HD cells exhibits a significant increase in the expression of ER stress markers genes (ATF4, GRP78, PERK, sXBPI) compared to control cells, indicating elevated proteostatic burden in HD condition. Reduced expression of ER stress marker genes confirms reduced ER stress upon VD treatment (n=3). Bar plot is from three independent experiments (n = 3) and all values are presented as mean with SD; *p value calculated using unpaired Welch’s t-test Holm’s correction *P < 0.05. *p < 0.05, **p < 0.005, ***p < 0.0005, and ****p < 0.00005).

## 3. Discussion

Huntington’s disease (HD) is characterized by progressive neuronal dysfunction, mitochondrial impairment, oxidative stress, and proteostasis defects^67,68^. Although vitamin D has been implicated in neuroprotection and mitochondrial regulation in several neurological contexts, its mechanistic contribution towards beneficial effects on cognitive impairment in HD remains obscure. In the present study, using both a chemical (3-nitropropionic acid; 3-NP) mouse model of HD and a cellular genetic model, we systematically dissect the regulation of the VD metabolic pathway, the impairment of which is linked to behavioral and organellar defects observed in HD pathology. VD supplementation improves behavioral performance and cellular viability by restoring mitochondrial and endoplasmic reticulum (ER) homeostasis, and rescues the expression of key genes involved in VD metabolism, including CYP27A1 and the VDR. Histopathological analysis of the striatum and hippocampus revealed neurodegenerative alterations, confirming the establishment of disease-associated pathology in the 3-NP mouse model and its recovery following VD supplementation, accompanied by improved CYP27A1 expression. In addition, VD supplementation attenuated ER stress markers, indicating improved proteostasis.

Furthermore, overexpression of mitochondrial CYP27A1 in the *in vitro* genetic model not only restored mitochondrial function but also enhanced VDR expression, promoting mitochondrial biogenesis and improving cellular viability under HD conditions. Collectively, these findings highlight a critical role for VD signaling in preserving mitochondrial and cellular homeostasis in HD, with CYP27A1 emerging as a key mediator of this protective mechanism and position the CYP27A1–VDR axis as a potential therapeutic target (Figure 8).

**Figure 8:**
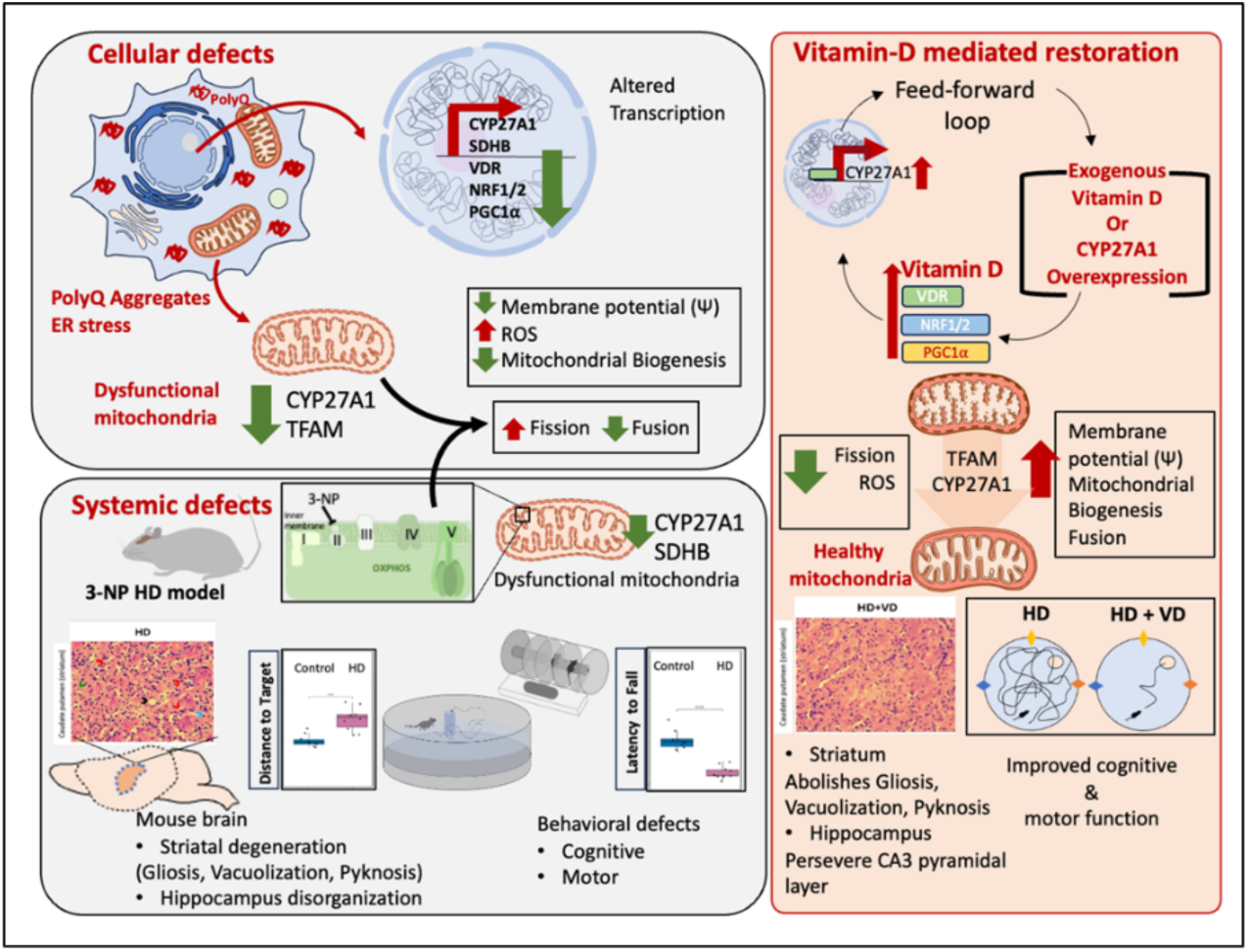
Schematic overview of the CYP27A1-VDR intracrine axis mediating organelle Homeostasis and cognitive rescue in Huntington’’s disease models. Graphical abstract summarizing the role of the CYP27A1-VDR axis in rescuing bioenergetic, proteostasis, and cognitive decline in Huntington’’s disease. Intracellular PolyQ aggregates and chemical complex Il inhibition (via 3-NP) induce severe cellular and systemic defects characterized by the down-regulation of mitochondrial CYP27A1, SDHB, and TFAM, accompanied by unbalanced mitochondrial fission, oxidative stress, and attenuated biogenesis pathways. At the tissue and organismal levels, these defects lead to striatal and hippocampal neurodegeneration, yielding profound motor deficits and spatial memory deterioration. Conversely, treatment via exogenous Vitamin D or direct CYP27A1 overexpression drives an intracrine positive feed-forward loop that upregulates VOR, NRF1/2, and PGCla. This signaling cascade restores complex Il expression, normalizes mitochondrial structural dynamics, and reduces ROS accumulation. Ultimately, this localized mitochondrial rescue protects the structural integrity of the hippocampal-striatal network, resolving histopathological lesions with subsequent rescue of cognitive and motor phenotypes.

### Behavioral Rescue: restoring Cognitive and Motor Function

A central feature of HD pathology is the development of motor and cognitive deficits associated with mitochondrial dysfunction. In particular, impaired expression and activity of mitochondrial Complex II represents a pathogenic defect that was effectively recapitulated in the 3-nitropropionic acid (3-NP) chemical model of HD^26^. A key finding of our study is the therapeutic efficacy of VD) in reversing behavioral deficits associated with HD. VD supplementation significantly improved cognitive and motor performance (**Supplementary Figure1B**). In 3-NP-induced HD mice, spatial memory was severely compromised, as evidenced by significantly increased escape latencies in the Morris Water Maze (Figure 1B). Remarkably, VD supplementation markedly mitigated these cognitive impairments, dramatically shortening escape latencies and demonstrating enhanced spatial learning and memory retention (Figure 1B, Supplementary Figure 1A).

This behavioral and functional recovery was supported by histopathological analysis indicating VD mediated neuroprotection. Histopathological evaluation revealed attenuation of vacuolization, gliosis, and neuronal loss in both the striatum and hippocampal CA3 region (Figure 1C). within both the striatum and the hippocampal CA3 region. By safeguarding these critical brain regions, VD preserves the structural integrity of the complex neural circuits that orchestrate cognition and motor coordination. To our knowledge, this study provides the first definitive evidence demonstrating that VD can successfully rescue cognitive deficits in an experimental model of Huntington’s disease, highlighting its potential as a disease-modifying neuroprotective agent.

### Molecular basis of rescue: Restoration of Mitochondrial Health

The observed behavioral improvements are supported by molecular evidence demonstrating that VD targets core bioenergetic deficits in HD. Consistent with clinical post-mortem studies and several prior reports, HD pathology involves impaired oxidative phosphorylation systems, particularly at mitochondrial Complex II. In our experimental framework, 3-NP–induced Complex II dysfunction—evidenced by a marked reduction in the SDHB within the striatum and hippocampus was effectively rescued by VD supplementation (**Figure 2A**). This restoration provides a mechanistic basis for improved mitochondrial respiration and behavioral recovery.

Beyond bioenergetic capacity, the structural dynamics of the mitochondrial network are pivotal to cell survival. Both 3-NP mice and Q23/Q74 cellular models exhibited disrupted mitochondrial dynamics, characterized by downregulation of fusion markers (MFN1/2) and upregulation of fission proteins (DRP1, MFF), a phenotype tightly aligned with mutant huntingtin (mHtt)-mediated activation of DRP^70,72^ **(Figure 3A, Supplementary 3D**). VD supplementation corrected this imbalance by enhancing fusion and suppressing fission, thereby restoring mitochondrial network integrity. This rebalancing stabilized membrane potential, reduced oxidative stress, and improved cellular viability, ultimately preventing neuronal loss associated with HD.

### Transcriptional Renewal: The VDR–PGC-1α–NRF1–TFAM Biogenesis Cascade

Mitochondrial biogenesis fundamentally relies on the coordinated regulation of nuclear and mitochondrial gene expression, a process orchestrating a complex transcriptional network. Central to this regulatory network Nuclear Respiratory Factor 1 (NRF1), is known to directly control the transcription of Mitochondrial Transcription Factor A (TFAM) gene via binding to its promoter, thereby driving mitochondrial DNA (mtDNA) replication and translation. This critical NRF1–TFAM axis is classically co-activated by Peroxisome proliferator-activated receptor gamma coactivator 1-alpha (PGC-1α) to fuel mitochondrial turnover and maintain oxidative metabolism. In the context of HD, however, impaired PGC-1α expression and function have been widely implicated in the breakdown of mitochondrial biogenesis and consequent bioenergetic failure^52,69,71^. Crucially, literature shows that PGC-1α can physically and functionally serve as a potent transcriptional coactivator of the VDR itself^57^. This structural intersection provides a direct molecular link connecting VD-dependent transcriptional activity to mitochondrial gene regulation. We have observed significant increase of the expression of these transcriptional regulators (NRF-1, PGC-1α, TFAM, VDR) after VD supplementation in HD condition (**Figure 5A**). In perfect agreement with the enhanced expression of these upstream transcription factors, our data reveal that VD treatment also significantly improved mtDNA copy numbers (**Figure 5 B**) This quantitative increase confirms the activation of functional mitochondrial biogenesis, demonstrating that VD does not merely preserve existing, damaged organelles, but actively drives the synthesis of a fresh, bioenergetically competent mitochondrial pool to rescue compromised cells from metabolic failure. Overall, our findings suggest that rescuing VDR availability via VD supplementation potentially repairs the compromised VDR–PGC-1α interaction. By reinforcing expression of these transcriptional factors, VD restarts the downstream NRF1–TFAM signaling cascade, providing a powerful transcriptional mechanism to counteract HD-induced bioenergetic decline.

### The VD–CYP27A1–VDR Axis: A bioenergetic rheostat in HD controlling mitochondrial fission-fusion balance

To understand how VD restores mitochondrial health in HD, we dissected the VD metabolic pathway and identified a previously unrecognized VD–CYP27A1–VDR regulatory feedback loop that is markedly disrupted under disease conditions. Importantly, the brain maintains an intrinsic VD metabolic system through enzymes such as CYP27A1, CYP27B1, CYP2R1, and CYP24A1, enabling local (intracrine) regulation of VDR signaling independent of systemic VD levels^40,73^. CYP27A1 is expressed in neural tissue and contributes to sterol metabolism, thereby linking lipid homeostasis with mitochondrial function^74^.This local regulatory system becomes particularly important in pathological conditions were circulating VD availability may not adequately meet neuronal demand^75^

Although mutant huntingtin (mHtt) is the primary driver of HD pathology, the downstream metabolic vulnerabilities associated with disease progression remain incompletely understood. Our data reveal that HD conditions significantly downregulate CYP27A1, the principal mitochondrial enzyme responsible for 25-hydroxylation of VD, suggesting the existence of a localized metabolic deficit that limits the generation of active VD metabolites and consequently exacerbates mitochondrial dysfunction. Notably, VD supplementation reversed this deficit by restoring CYP27A1 expression in HD cells (**Figure 4B-C**).

Given that reduced CYP27A1 expression appeared closely associated with impaired mitochondrial function and altered VDR-mediated signaling, we hypothesized that targeted CYP27A1 overexpression would phenocopy the protective effects of VD treatment in *in vitro* HD models. Consistent with this hypothesis, CYP27A1 overexpression alone restored the expression of key mitochondrial regulatory transcription factors, including PGC-1α, TFAM, and VDR, while simultaneously improving mitochondrial and cellular health **(Figure 6A-B, Supplementary Figure 6A**). This was accompanied by enhanced MFN1 expression, reduced mitochondrial fission signatures, and significantly decreased apoptosis in HD cells (**Figure 6D**), collectively supporting a direct role for CYP27A1 in maintaining mitochondrial homeostasis.

To further investigate the mechanistic basis of this regulation, we analyzed publicly available VDR ChIP-seq datasets and identified significant VDR occupancy at the CYP27A1 promoter region (**Figure 6C**), indicating direct transcriptional regulation of CYP27A1 by VDR. These findings position CYP27A1 both upstream and downstream of VDR signaling within the VD metabolic pathway, thereby establishing a feed-forward regulatory loop capable of amplifying mitochondrial protective responses and restoring mitochondrial biogenesis and dynamics.

In addition to its canonical nuclear functions, emerging evidence indicates that VDR can localize to mitochondria and directly regulate mitochondrial gene expression through interactions with mitochondrial nucleoids and TFAM^48,76,77^. Moreover, PGC-1α has been shown to function as a coactivator of VDR, further enhancing its transcriptional activity^57^. Consistent with these observations, we detected coordinated activation of the VDR–PGC-1α–NRF1–TFAM signaling axis, which was associated with increased mtDNA content and synchronized upregulation of both nuclear- and mitochondrial-encoded respiratory genes. Together, these findings support a model in which VD restores mitochondrial integrity in HD through a tightly interconnected CYP27A1–VDR regulatory circuit that reinforces mitochondrial biogenesis, dynamics, and cellular survival.

### Regulating organellar (mitochondria and ER) proteostasis is linked with normalizing the CHCHD4 Compensatory Response

During proteotoxic stress, upregulation of the mitochondrial import factor Mia40 (CHCHD_4_ in humans) is required to maintain the disulfide relay system and keep essential protein import flowing. Earlier work by Schlagowski et al. demonstrated that they can prevent polyQ aggregation by bolstering cytosolic proteostasis ^36^. We identified a critical angle involving this protein in our HD models. We observed that CHCHD4 levels were significantly elevated in both 3-NP mice and HD cells, likely as a dysfunctional compensatory mechanism to cope with cellular proteotoxic stress influenced by the presence of cytosolic mHtt aggregates and dysfunctional mitochondria (**Figure 3E**). In our HD cellular model, VD treatment significantly reduced the burden of GFP-positive mHtt aggregates in insoluble protein fractions (**Supplementary Figure 7**). This reduction in aggregate load was accompanied by marked attenuation of the ER-specific Unfolded Protein Response (UPR^ER^), as indicated by decreases in key ER stress markers Spliced XBP1, GRP78, and PERK–ATF4 (**Figure 7**). Overall, VD supplementation does not simply suppress these proteotoxic stress; rather, it normalizes CHCHD4 expression. VD supplementation did not further increase CHCHD4; instead, it normalized its expression. By restoring mitochondrial dynamics (fusion) and the CYP27A1–VDR–PGC-1α transcriptional axis, VD provides the cell with sufficient bioenergetic stability (improved membrane potential and reduced ROS) to resolve the underlying stress **(Figure 3C-D**). Thus, VD eliminates the need for this desperate compensatory response, leading to a more robust and stable cellular proteostasis environment. Additionally, by modulating the fission-fusion equilibrium and reinforcing the mitochondrial network, VD acts as a master coordinator of cellular health by improving cellular viability (Figure 2F).

In conclusion, this study establishes the CYP27A1–VDR axis as a master regulator of neuroprotection and organellar homeostasis in Huntington’s disease, demonstrating for the first time that Vitamin D (VD) supplementation can successfully rescue severe cognitive deficits in experimental models of HD. By integrating pharmacological and genetic models, we demonstrate that vitamin D (VD) supplementation is not merely a supportive intervention but a potent molecular strategy that targets the core bioenergetic and proteostatic failures of HD.

By repairing a locally compromised, intracrine feed-forward loop VD treatment acts as a bioenergetic rheostat that suppresses toxic mitochondrial fission and drives functional mitochondrial biogenesis via the VDR–PGC1α–NRF1–TFAM transcriptional cascade. This comprehensive restoration of the mitochondrial network effectively bridges organellar crosstalk, clearing mutant huntingtin aggregate load, silencing ER stress pathways, and uniquely normalizing the mitochondrial CHCHD4 compensatory response to alleviate the global proteostatic burden and balance cellular fate during disease stress.

While these findings shift the therapeutic paradigm from passive organellar preservation to active renewal, certain limitations warrant consideration: the 3-NP chemical model mimics acute Complex II inhibition rather than the progressive, full-length HTT genetic context of HD, necessitating future validation in transgenic models; the exact proportional contribution of local brain-derived VD metabolism versus systemic metabolites remains to be fully delineated *in vivo*; and translating these experimental doses into safe, hypercalcemia-free human regimens requires careful calibration. Despite these constraints, this integrated rescue mitigates accelerated neuronal death and positions the local CYP27A1–VDR signaling circuit as a high-value, druggable therapeutic target to counteract metabolic failure and preserve long-term neural circuit integrity in Huntington’s disease.

## 4. Methodology

### 4.1 Cell Culture Model and Experimental Design for *In Vitro* HD Study

Recombinant HEK293 cell lines engineered to express exon 1 of the human HTT gene with either 23 CAG repeats (control line-Q23) or 74 CAG repeats (HD line-Q23), fused to GFP under the control of a doxycycline-inducible promoter, obtained as a generous gift from David C Rubinsztein’s lab, University of Cambridge, UK. Both the cell lines were tested for mycoplasma contamination using the EZdetect PCR kit and were confirmed to be negative (HiMedia Laboratories Private Limited). Dulbecco’s Modified Eagle’s Medium (DMEM, Gibco, Thermo Fisher Scientific, USA) supplemented with 10% heat-inactivated fetal bovine serum (Gibco, Thermo Fisher Scientific) and an antibiotic cocktail [10000 units/ml penicillin, 10 mg/ml streptomycin (Gibco, Thermo Fisher Scientific), Hygromycin (150 µg/ml), blasticidin (5 µg/ml)] was used to culture Q23 and Q74 cell lines in poly-L-ornithine coated plates. The cells were maintained at 37C in 5% Co2 atmosphere. At 70% confluence, polyQ HTT formation was induced by adding 1ug/ml doxycycline for 48 hours. To assess VDR expression, other mitochondrial gene levels, and the effects on mitochondrial morphology and cell viability, the cell lines were treated with, 1α,25-dihydroxyvitamin D₃ (Calcitriol; 50nM, Sigma Aldrich) or Calcitriol (50nM, HY10002, MedChemExpress) for 48 hours, as depicted in the schematic. To maintain the consistency, for all cell culture experiments, Calcitriol treatment is referred as Vitamin D (VD) treatment or supplementation.

### 4.2 Transient Overexpression of CYP27A1 in Mammalian Cells

Cells were seeded one day prior to transfection to achieve ∼60–80% confluency at the time of transfection. Transient overexpression was performed using the pcDNA3.1(+)-CYP27A1 construct or empty vector as control. For each well of a 6-well plate, 2 µg plasmid DNA was diluted in 200 µL jetPRIME buffer (Polyplus Transfection) and mixed gently. Subsequently, 4 µL jetPRIME reagent was added, and the mixture was vortexed briefly followed by incubation at room temperature for 10 minutes to allow formation of transfection complexes. After 4–5 hours, the medium was replaced with an antibiotic selection medium. Cells were then incubated for 48 hours, after which they were harvested for downstream assays.

### 4.3 Animal model and experimental design

All animal experiments were conducted in accordance with institutional ethical guidelines and approved by the Institutional Animal Ethics Committee. Male C57BL/6 mice (12-13 weeks old; ∼26 ± 3 g) were procured and housed under standard laboratory conditions with access to food and water, maintained at 25 ± 2°C under a 12 h light/dark cycle. Animals were acclimatized for 5 days prior to experimentation. To model Huntington’s disease (HD), a 3-nitropropionic acid (3-NP)-induced neurodegeneration model was employed. Mice were randomly divided into four experimental groups: (i) Control (saline), (ii) HD (3-NP), (iii) Vitamin D (VD), and (iv) HD + VD. HD-like pathology was induced by intraperitoneal (i.p.) administration of 3-NP at 25 mg/kg, given in three doses at 12 h intervals (cumulative dose: 75 mg/kg), as previously described in Manjari, et al., and control animals received equivalent volumes of saline ^10^. Vitamin D_3_ (cholecalciferol) was administered i.p. at 500 IU/kg/day for 15 consecutive days. In the HD + VD group, VD treatment was initiated 24 h after the final 3-NP injection and continued daily for 15 days. To maintain the consistency, for all mouse experiments, vitamin D_3_ (cholecalciferol) treatment is referred as Vitamin D (VD) treatment or supplementation.

### 4.4 Behavioral Assessment and Brain Tissue Collection

Spatial learning and memory were assessed using the Morris water maze (MWM) paradigm, performed as described previously (Morris, 1984; Vorhees and Williams, 2006)^78^. Mice were tested in a circular water tank filled with opaque water, surrounded by four distinct visual cues placed around the testing arena. A hidden escape platform was submerged below the water surface in one quadrant. Animals were trained over 5 consecutive days, during which each mouse was allowed to locate the hidden platform using spatial cues. This was followed by testing sessions over three days, and finally a probe trial, in which the platform was removed to assess memory retention. The latency to reach the platform, speed at which the mice swim, and many other parameters were recorded. During the probe trial, the time spent in the target quadrant and search strategies were quantified. Swimming trajectories were recorded and analyzed using tracking software from Panlab - SMART video tracking system (Panlab Harvard Apparatus, Spain). Following completion of behavioural assessments, animals were sacrificed on the experimental endpoint. Mice were anesthetized using isoflurane and immediately decapitated. The striatum and hippocampus were carefully isolated and processed for downstream molecular and biochemical analyses.

### 4.5. Histopathological (Hematoxylin and Eosin staining) and Immunohistochemistry analysis

Following completion of behavioral assessments, mice were deeply anesthetized using isoflurane and subjected to transcardial perfusion. Briefly, the thoracic cavity was opened to expose the heart, and a perfusion needle was inserted into the left ventricle, while the right atrium was incised to allow drainage. Animals were first perfused with ice-cold phosphate-buffered saline (PBS) to remove circulating blood, followed by perfusion with 4% paraformaldehyde (PFA) for fixation. Successful perfusion was confirmed by clearing of the liver and extremities ^79^. Following perfusion, brains were carefully dissected and post-fixed in 4% PFA at 4°C, and subsequently processed for histological and immunohistochemistry (IHC) analysis (**Supplementary File 1**).

#### H and E staining

Paraffin-embedded tissues were sectioned at 4–5 µm thickness using a rotary microtome and mounted onto glass slides. Sections were deparaffinized in xylene, rehydrated through descending grades of ethanol, and stained with hematoxylin to visualize nuclei, followed by eosin staining to visualize cytoplasmic and extracellular components. Stained sections were dehydrated, cleared, and mounted using DPX mounting medium. Histological evaluation was performed using a light microscope at 20× magnification, focusing on the caudate putamen (striatum) and hippocampal subregions (DG, CA3, and CA1).

#### IHC Staining

Paraffin-embedded tissue sections were heated at 55–60°C for 10–15 min, deparaffinized in three changes of xylene for 5 min each, and rehydrated through graded alcohols. Sections were incubated in three changes of 100% ethanol for 5–8 min each, followed by 90% and 80% ethanol for 6–8 min each, rinsed in distilled water, and washed with PBS. Endogenous peroxidase activity was blocked using 1–3% hydrogen peroxide prepared in methanol, followed by washes with distilled water and PBS.

Antigen retrieval was performed using preheated retrieval buffer in a microwave for three cycles of 5 min each, with 1 min intervals between cycles. For skin tissue, Tris-EDTA buffer, pH 9.0, containing 10 mM Tris base, 1 mM EDTA, and 0.05% Tween-20 was used. Slides were allowed to cool, rinsed with distilled water, and washed with PBS. Sections were blocked with 0.25% BSA for 30 min at room temperature and incubated with the primary antibody at the appropriate dilution for 1 h at room temperature or overnight at 4°C. After washing twice with PBS for 4 min each, sections were incubated with the appropriate secondary antibody for 60 min at room temperature. Immunoreactivity was developed using ready-to-use DAB substrate containing DAB chromogen for approximately 5 min. Slides were washed in running tap water, counterstained with hematoxylin for 3 min, differentiated with 1% acid alcohol, dehydrated through 80%, 90%, and 100% ethanol, cleared in three changes of xylene, and mounted using DPX.

### 4.6 Quantitative RT-PCR

Total RNA was extracted from control and HD cell lines using the Trizol method (RNAiso Plus, Takara) following 48 hours of treatment. RNA quantity and quality were determined using a NanoDrop spectrophotometer (Thermo Scientific, USA). cDNA was prepared by reverse transcribing 1 µg of total RNA with the PrimeScript RT reagent kit (Takara).

RT-qPCR was performed in 96-well plates on either the LightCycler 480 (Roche) or CFX96 Real-Time PCR system (Bio-Rad) with SYBR green master mix (TB Green, Takara) with gene-specific primers (**Supplementary File 2**). PCR conditions used are an initial 95°C for 10 min, followed by 45 cycles of 95°C for 15 s, 60°C for 10 s, and 72°C for 10 s. Each gene was analyzed in three technical replicates across three independent experiments. Relative expression was calculated using the ΔCt method with β-actin as the reference gene, where ΔCt = Ct(target) - Ct(β-actin). Fold changes between treated and control groups were determined by the ΔΔCt method, where fold change = 2−(−ΔΔCt). Here, ΔCt (cycle difference) = Ct (target gene) - Ct (control gene), and ΔΔCt = ΔCt (treated condition) - ΔCt (control condition).

### 4.7 Confocal microscopy

Confocal dishes were seeded with control HD cell lines at a density of 1.8 × 10−6 cells and maintained at 37°C in a 5% CO₂ incubator. Once cultures reached approximately 60–70% confluence, expression of polyQ HTT was induced with 1 µg/ml doxycycline (Sigma-Aldrich) for 6 hours, after which cells were treated with Vit-D3 (50 nM, Sigma-Aldrich) for an additional 48 hours prior to imaging. Before acquisition, dishes were placed in a temperature- and CO₂-controlled chamber mounted on a Leica SP8/DMi8 confocal microscope. Cells were incubated with MitoTracker Red (100 nM, Invitrogen, Thermo Scientific) for 5 minutes at 37°C, and images were collected using the appropriate excitation/emission settings (581 nm/644 nm) for MitoTracker and 475 nm/509 nm for GFP. Fluorescence and corresponding bright-field images were recorded from multiple fields using identical acquisition parameters.

Image processing was performed in Fiji (ImageJ, NIH), where brightness and contrast were adjusted using the standard “Brightness/Contrast” tool and basic LUT modifications to optimize visualization without altering the underlying data. Quantitative analysis of mitochondrial morphology was carried out using the MiNA plugin, following the developer’s guidelines (Stuart Lab, https://github.com/stuartlab).

### 4.8 Western blotting

Control(Q23) and HD(Q74) cell lines were collected after 48 hours of doxycycline induction and Vit-D3 treatment, lysed in RIPA buffer by continuous agitation for 1 minute and left on ice for 5 to 10 minutes. The cell lysate was centrifuged at 16.000xg for 20 min at 4°C and the supernatant containing total protein was collected. The concentration of the protein was measured using a BCA protein assay kit (Pierce, Thermo Fisher). Total proteins (75µg) were resolved by SDS-PAGE and subsequently transferred to a polyvinylidene fluoride (PVDF) membrane (Immobilon PVDF, IPVH00010, Millipore). The membrane was blocked in 8% bovine serum albumin (BSA) for one hour, followed by primary antibody incubation overnight at 4℃ with agitation. The following antibodies were used: VDR (A23289, 1:1000, ABclonal), DRP1 (A21968, 1:1000, ABclonal), Mitofusin-1/MFN1 (A21293, 1:1000, ABclonal), Mitofusin-2/MFN2 (D2D10, 1:1000, CST), SDHB (A10821, 1:1000, ABclonal), β-Tubulin (AC008, 1:1000, ABclonal), FIS1 (E3K90, 1:1000, CST), SDHA (A13852, 1:1000, Abclonal), MFN1 (A21293, 1:1000, Abclonal), CHCHD4 (A25029, 1:1000, Abclonal), COX4 (A11631, 1:1000, Abclonal), TAFM (A3173, 1:1000, Abclonal). The next day, after washing the blots with Tris-buffered saline with 0.1% Tween 20 detergent (TBST), secondary antibody was added and incubated for one hour at room temperature with agitation. The antibody used was: Anti-rabbit IgG, HRP-linked antibody (1:10,000, 7074S, CST). The signal intensities of the bands were captured using the fusion pulse gel documentation system (Eppendorf, USA). ImageJ software was used to quantify the band intensities. Using Image J software, the protein expression was measured and normalized to the protein intensity of housekeeping gene **(Supplementary File 3**)

### 4.9 Determination of mitochondrial membrane potential and ROS

For mitochondrial ROS and membrane potential analysis, control and HD cells were plated in 6-well plates and incubated for 24 hours under optimal growth conditions. To induce PolyQ expression, doxycycline was added to the medium at a final concentration of 1 μg/mL, and the cells were further incubated for 6 hours. After induction, cells were supplemented with Vit-D3 at 50nM concentration and incubated for an additional 48 hours before analysing in flow-cytometer.

After 48 hours, cells were trypsinized and washed with PBS, and resuspended in PBS containing 5 μM MitoSOX Red (M36008, Invitrogen). This suspension was incubated for 15 minutes at 37 °C in the dark to allow for MitoSOX Red uptake. Fluorescence acquisition was performed using a BD FACS Aria flow cytometer with excitation and emission wavelengths of 488 nm and 585 nm, respectively. Mitochondrial ROS generation in treated and control cells was analysed. Fluorescence intensity values from 10,000 cells were recorded and visualized in FlowJo software. Statistical significance of ROS changes was determined from the median fluorescence intensity across three independent biological replicates (n = 3).

To evaluate mitochondrial membrane potential (ΔΨm), cells were stained with 100 nM TMRE for 20 minutes at 37 °C. For the positive control, FCCP (10 μM) was added to the cells 1 hour prior to TMRE staining to induce mitochondrial depolarization. After staining, cells were trypsinized, washed twice with PBS, centrifuged, and resuspended in PBS. Samples from different treatment groups were then analyzed using a BD FACS Aria flow cytometer to quantify TMRE fluorescence intensity and assess changes in mitochondrial membrane potential. TMRE fluorescence from 10,000 cells per condition was analyzed using FlowJo software to visualize alterations in membrane potential. Statistical comparisons were made using the median fluorescence intensity from three independent biological experiments (n = 3).

### 4.10 Apoptosis Assay

Control and HD cells were plated in culture plates and allowed to adhere for 24 hours under standard growth conditions. Further, doxycycline was added at a final concentration of 1 μg/mL to induce PolyQ expression, and incubation continued for an additional 6 hours. Subsequently, cells were treated either with Vit-D3 at 50nM or overexpressed with CYP27A1. The treated cells were then maintained for 48 hours before proceeding to apoptosis analysis (**Supplementary File 4**).

Apoptosis was evaluated using the Annexin V-Elab Fluor Red 780/Propidium Iodide (PI) assay (E-CK A23, Elab Bioscience, Texas, USA) according to the manufacturer’s instructions. Briefly, cells were harvested, washed twice with PBS, and resuspended in 100 μL of 1× Annexin V binding buffer (1 × 10⁵ cells per sample). After centrifugation at 300 × g for 5 min, supernatants were discarded, and the pellets were washed again with PBS to remove debris. The cells were gently resuspended and counted to ensure uniform sample distribution.

Each tube containing 1 × 10⁵ cells was stained with 2.5 μL Annexin V-Elab Fluor Red 780 and 2.5 μL PI, followed by incubation for 15 min at room temperature in the dark. After staining, 400 μL of 1× binding buffer was added, and the samples were gently mixed before analysis. Flow cytometric acquisition was carried out immediately using a BD FACS Aria system, detecting Annexin V-Elab Fluor Red 780 (625/765 nm) and PI (488/610 nm) fluorescence. A total of 10,000 cells were analyzed per sample, and data were represented as dot plots using BD acquisition and analysis software to assess the extent of apoptosis at various conditions.

### 4.11 VDR ChIP-seq analysis

VDR ChIP (SRR26136086) and input control (SRR26136085) SRA files were downloaded from the NCBI SRA database using the prefetch v3.2.1 command of the SRA Toolkit and converted to fastq format using fastq-dump. The reads were trimmed using Trimmomatic v0.39 with the parameters “TruSeq3-SE.fa:2:30:10 LEADING:3 TRAILING:3 SLIDINGWINDOW:4:15 MINLEN:36”. The quality of the raw and trimmed reads was assessed using FastQC v0.12.1. The trimmed reads of each sample were aligned to the human reference genome GRCh38 using HISAT2 (v2.2.1) with default parameters. The resultant BAM files were processed using samtools v1.21 to remove duplicates, and exclude reads mapped to uncharacterized chromosomes, chrM, and chrY, as well as the ENCODE blacklist regions (https://www.encodeproject.org/files/ENCFF356LFX/@@download/ENCFF356LFX.bed.gz).

The processed BAM files were used to call peaks using MACS2 v2.2.9.1 with default parameters (**Supplementary File 5**).

### 4.12 Chromatin Immunoprecipitation (ChIP)-PCR

ChIP assays were carried out using the SimpleChIP ® Enzymatic Chromatin IP Kit (Cell Signaling Technology, Cat No: 9003S) according to manufacturer’s instructions. Briefly, control/HD cell lines were cultured for 48 h in 10 cm dishes until ∼90% confluency. Further, cross-linking was performed using 1% formaldehyde by directly adding it to the culture medium and incubating for 10 minutes at 37 °C. Cross-linking was quenched by adding 0.125 M glycine for 5 min at room temperature. Cells were then washed twice with ice-cold PBS and harvested by scraping into PBS containing protease inhibitors, followed by centrifugation at 2000 × g for 5 min at 4 °C. The resulting cell pellets were either processed immediately or stored at –80 °C.

For nuclear isolation, pellets were resuspended in Buffer A supplemented with protease inhibitors and DTT, incubated on ice for 10 minutes, and centrifuged to collect the nuclei. Nuclei were washed twice with Buffer B and resuspended in 1X ChIP Buffer. Chromatin was digested with micrococcal nuclease at 37 °C for 20 min to yield fragments of ∼150–900 bp. Reactions were stopped with EDTA, and nuclei were lysed by homogenization. The lysates were clarified by centrifugation, and supernatants containing cross-linked chromatin were stored at – 80 °C until use.

For immunoprecipitation, 5–10 µg of digested chromatin was diluted in ChIP buffer and incubated overnight at 4 °C with the anti-VDR antibody. Normal rabbit IgG was used as a negative control. Protein G magnetic beads were then added and incubated for 2 hr at 4 °C with rotation. Beads were then washed three times with a low-salt wash buffer and once with a high-salt wash buffer. Chromatin was eluted from beads in ChIP elution buffer for 30 min at 65 °C. Cross-links were reversed by incubation at 65 °C for 2 hr in the presence of NaCl and Proteinase K. DNA was then purified using spin columns according to the manufacturer’s instructions. Input and ChIP DNA were analyzed by PCR with primers specific for RPL30 as positive control and CYP27A1 the VDR target gene, to confirm VDR binding enrichment (Supplementary File 6).

### 4.13 Statistical analysis

Post-processing data analysis and statistical analyses were done in R version 4.3.3 (R Core Team, www.Rproject.org) using specific packages. The CRAN R packages: “tidyverse” and “devtools” were used for basic data manipulation, visualization and statistical analysis. These packages allowed access to ‘rstatix’ (statistical calculations), ‘ggpubr’, ‘ggplot2’, ‘dplyr’, ‘tidyr’, ‘readr’, ‘purrr’, and ‘tibble’ libraries for data analysis and visualization. P-Value was calculated through pairwise t-test with default Bonferroni correction indicated in the respective figure legends. The p-value significance is denoted by an asterisk (*) (* represents the following p-values: *p < 0.05, **p < 0.005, ***p < 0.0005, ****p < 0.00005).

## Authors’ contributions

S.M. conceptualized and supervised the project. V.K. performed all the wet-lab experiments and mice behavioral studies. P.K. (Pradeep Kodam), KC and SKVM helped in mice work, P.K. (Pradeep Kodam) helped in cloning, survivability assay and ChIP-qPCR. SNB helped in western blot experiments. GB and AS analysed ChIP-Seq data. P.K. (Pragya Komal) performed brain dissection. S.P. and S.C. helped V.K. in performing histopathology of the brain samples. S.M. & V.K. wrote the original draft. A.S, P.K (Pragya Komal), & S.C. helped with critical assessment and edits. All authors have read and agreed to the final version of the manuscript.

## Acknowledgements

We thank David C Rubinsztein’s lab, University of Cambridge, UK for providing Recombinant HEK293 cell lines (containing exon 1 of the human HTT gene, with either 23 CAG (control; Q23) or 74 CAG (HD; Q74) repeats, fused to Green Fluorescent Protein (GFP). We thank Dr. Victoria Barratt from David Rubinsztein Group for helping us to get the cell lines and sharing the details of growth conditions. The contribution of Central Analytical Laboratory (CAL), Birla Institute of Technology and Science-Pilani (BITS-Pilani), Hyderabad Campus, is gratefully acknowledged for providing access to BD FACS Aria II, Leica confocal microscope (Leica SP8/DMi8), RT-PCR system (BIORAD CFX Opus 96). We also acknowledge the DDT-FIST (SR/FST/LS1-526/2012) facility of the Department of Biological Sciences, BITS-Pilani, Hyderabad campus for providing access to RT-PCR (LC480 Roche, Roche), imaging system for Western blot (Fusion Pulse 6, Vilber Lourmat). DBT Builder (BT/INF/22/SP42551/2021) for providing mouse facility to conduct behavioral tests as well as brain dissection. S.M. acknowledges ICMR Ad-hoc research grants (Project ID 2021-13896 and 2021-13256), and BITS-Pilani (Hyderabad Campus) for OPERA (OPERA-1022) award. V.K., K.C. SNB, and SKVM acknowledged BITS-Pilani (Hyderabad Campus) for institutional doctoral fellowship and P.K. (Pradeep Kodam) acknowledges CSIR/UGC/NFOBC for doctoral fellowship. K.C. also acknowledges SERB-SRG for the JRF fellowship.

